# PP2C phosphatases promote autophagy by dephosphorylation of the Atg1 complex

**DOI:** 10.1101/309500

**Authors:** Gonen Memisoglu, Vinay V. Eapen, James E. Haber

**Affiliations:** Department of Biology and Rosenstiel Basic Medical Sciences Research Center, Brandeis University, Waltham, MA 02454, USA

## Abstract

Autophagy is orchestrated by the Atg1-Atg13 complex in budding yeast. Under nutrient-rich conditions, Atg13 is maintained in a hyperphosphorylated state by TORC1 kinase. After nutrient starvation, Atg13 is dephosphorylated, triggering Atg1 kinase activity and autophagy induction. The phosphatases that dephosphorylate Atg13 remain uncharacterized. We show that two redundant PP2C phosphatases, Ptc2 and Ptc3, regulate autophagy via dephosphorylating both Atg13 and Atg1. In the absence of these phosphatases, starvation-induced macroautophagy is inhibited, as is the cytoplasm-to-vacuole targeting (Cvt) pathway, and the recruitment of the essential autophagy machinery to phagophore assembly sites (PAS) is impaired. Despite prolongation of the DNA damage-induced checkpoint in *ptc2*Δ *ptc3*Δ cells, genotoxin-induced autophagy is also blocked. Creating a genomic *atg13-8SA* allele under its endogenous promoter, lacking key TORC1 phosphorylation sites, bypasses the autophagy defect in *ptc2*Δ *ptc3*Δ strains. Taken together, these results imply that PP2C type phosphatases promote autophagy by regulating Atg1 complex.

## INTRODUCTION

Macroautophagy is a catabolic recycling pathway that is responsible of degradation of cytoplasmic components by targeting them to the vacuole under nutrient-depleted conditions (Nakatogawa et al. 2009, Wen and Klionsky 2016). During the early steps of macroautophagy, phagophores are formed around cytoplasmic components (Suzuki et al. 2001) and nucleated around a phagophore assembly site (PAS) adjacent to vacuoles. Then, phagophores expand to form mature autophagosomes, which eventually fuse with the vacuole (Abeliovich et al. 2000), wherein the contents engulfed within the autophagosomes are degraded (Epple et al. 2001). In addition to the degradation of random portions of the cytoplasm for recycling purposes under nutrient-depleted conditions, specific organelles or molecules can be targeted for degradation or processing under nutrient-repleted conditions (Farre and Subramani 2016) via selective autophagy. Selective autophagy pathways rely on the recognition of cargo via specific receptor proteins and the scaffold protein, Atg11 in yeast (Kamber et al. 2015, Kim et al. 2001). One of the best-studied selective autophagy pathways is cytoplasm-to-vacuole (Cvt) pathway, which targets at least two precursor hydrolases, prApe1 and α-mannosidase, to the vacuole for their maturation through proteolytic cleavage (Lynch-Day and Klionsky 2010). The genetic requirements of Cvt and macroautophagy significantly overlap, however there are some proteins that specifically affect one of the two pathways. Notably, the scaffold protein Atg17 is required specifically for macroautophagy but is dispensable for Cvt (Kabeya et al. 2005, Kamada et al. 2000). Conversely, Atg19, the receptor protein for the Cvt cargo, and the scaffold protein Atg11 are important for Cvt function, but not for macroautophagy (Sidney V. Scott et al. 2001, Yorimitsu and Klionsky 2005).

The sensor of the availability of the nutrients is the target of rapamycin complex, TORC1. When nutrients are abundant, Atg13, an essential part of the core autophagy machinery, is kept in a hyperphosphorylated and inhibited state (Kamada et al. 2010). In addition to Atg13, the central kinase in the autophagy pathway, Atg1, is phosphorylated and inhibited by PKA and TORC1-dependent phosphorylations (Matsuura et al. 1997, Stephan et al. 2009, Stephan et al. 2010). When the nutrients are scarce, TORC1 activity is inhibited; consequently, Atg13 is rapidly dephosphorylated, this dephosphorylation of Atg13 is required for stimulation of Atg1’s kinase activity upon autophagy induction (Kamada et al. 2010). Recent work showed that Atg1 and Atg13 binding is constitutive; however, Atg1-Atg13 interaction is enhanced upon Atg13 dephosphorylation (Kraft et al. 2012, Yeh et al. 2011b). Atg1 and Atg13 also bind to the Atg17-Atg29-Atg31 scaffolding sub-complex to form a complete Atg1 kinase complex upon autophagy induction (Kabeya et al. 2009, Kawamata et al. 2008, Suzuki et al. 2007). The Atg1 complex localizes at the PAS and activates downstream *ATG* proteins to promote the formation of autophagosomes (Papinski et al. 2014).

The Atg1 complex is extensively regulated by several phosphorylations. Mass spectrometry studies collectively identified more than a dozen phospho-sites on Atg1 (Budovskaya et al. 2005, Papinski et al. 2014, Yeh et al. 2011a); however, only a few of these sites have been studied in depth *in vivo*. The autophosphorylation of Atg1-T226 that resides in the kinase activation loop is required for proper autophagy induction and Atg1’s kinase activity, yet is dispensable for Atg1’s binding to Atg13 (Kijanska et al. 2010, Yeh et al. 2010). Conversely, the phosphorylation of Atg1-S34 is inhibitory for its kinase activity and also for autophagy (Yeh et al. 2011a). Moreover, PKA-dependent phosphorylation of Atg1 on two serine residues inhibit its localization to the PAS (Budovskaya et al. 2005). As mentioned above, Atg13 is inhibited by TORC1-dependent phosphorylations on at least 8 serine residues (Kamada et al. 2010). Overexpression of *ATG13-8SA*, in which 8 serines were mutated to alanines, results in autophagy induction that is uncoupled from TORC1 activity even when there is a genomic WT copy of *ATG13* present. A recent study showed that the alanine substitution of 5 serine residues located in the MIM domain of Atg13 reduced Atg1-Atg13 binding and also impaired autophagy, suggesting that additional sites on Atg13 play a role in its regulation (Fujioka et al. 2014).

Although the post-translational regulation of the Atg1 kinase complex has been well-studied, the phosphatases that are responsible of the reversal of these phosphorylations remain uncharacterized. Previous work showed that PP2A-type phosphatases might be involved in Atg13 dephosphorylation in mammals (Wong et al. 2015). Furthermore, PP2A-type phosphatases are characterized as positive regulators of autophagy in mammals (Fujiwara et al. 2016) and in *Drosophila* (Banreti et al. 2012). In budding yeast, PP2A-type phosphatases have also been implicated in the dephosphorylation of Atg13 (Yeasmin et al. 2016); however, the role of PP2A-type phosphatases on autophagy still remains under debate, as they were reported to be negative regulators of autophagy by another group (Yorimitsu et al. 2009).

Previous work from our lab has shown that DNA damage induces a targeted, Atg11-dependent selective autophagy pathway, which we coined genotoxin-induced targeted autophagy (GTA) (Dotiwala et al. 2013, Eapen and Haber 2013, Eapen et al. 2017). GTA requires the core autophagy machinery, Atg1, Atg13, Atg14, Atg16 and Atg18 as well as the scaffold Atg11, in addition to the kinases that are required for the maintenance of DNA damage checkpoint, Mec1/ATR and Rad53/CHK2. A genome-wide screen to find modulators of GTA yielded Pph3, a PP4-type phosphatase as a negative regulator specifically of GTA but not of rapamycin-induced autophagy (Eapen et al. 2017). Pph3 has been demonstrated to dephosphorylate and inactivate the targets of checkpoint kinases, including the DNA damage-specific phosphorylated histone subunit, γ-H2AX and DNA damage checkpoint effector kinase Rad53 (Keogh et al. 2006, Kim et al. 2011, O’Neill et al. 2007, Sun et al. 2011). Consequently, deletion of PPH3 causes the cells to maintain a hyperactive DNA damage checkpoint.

Budding yeast’s PP2C-type phosphatases, Ptc2 and Ptc3, are serine/threonine-phosphatase paralogs that arose from whole genome duplication and are largely redundant (Maeda et al. 1993). They dephosphorylate a variety of targets including those involved in the high-osmolarity glycerol pathway (Warmka et al. 2001, Young et al. 2002), the unfolded protein response (Welihinda et al. 1998), the MAP-kinase pathway (Maeda et al. 1994) and the general stress response (Sharmin et al. 2014). In addition, these phosphatases counteract DNA damage checkpoint (DDC) signaling by directly dephosphorylating the central checkpoint kinase, Rad53 (Leroy et al. 2003). As a result, similar to the deletion of *PPH3*, deletion of *PTC2* and *PTC3* causes the hyperactivation of DDC in response to DNA damage (Kim et al. 2011, Leroy et al. 2003).

Since Ptc2 and Ptc3 play a comparable role as Pph3 for DNA damage checkpoint inactivation (Kim et al. 2011), we asked if Ptc2 and Ptc3 phosphatases have similar functions with Pph3 for GTA. Contrary to the suppression of GTA by Pph3, we find that Ptc2 and Ptc3 are positive regulators not only of GTA, but also of rapamycin-induced macroautophagy and Cvt. Deletion of *PTC2* and *PTC3* causes the accumulation of hyperphosphorylated Atg1 even in nutrient-replete conditions. Atg1 hyperphosphorylation is dependent on the kinase activity of Atg1, and on Atg13 and Atg11. In addition to Atg1, Atg13 is also maintained in a hyperphosphorylated state in the absence of *PTC2* and *PTC3*. The autophagy defect in the *ptc2Δ ptc3Δ* strain is suppressed by the constitutively active *ATG13-8SA* allele. Based on these results, we propose that the phosphatases Ptc2 and Ptc3 promote autophagy by dephosphorylating and stimulating Atg1 kinase complex.

## RESULTS

### PP2C-type phosphatases Ptc2 and Ptc3 promote DNA-damaged induced and rapamycin-induced autophagy

We used the GFP-Atg8 processing assay to monitor the autophagy induction in the phosphatase mutants (Klionsky et al. 2016). Upon autophagy induction, phosphatidylethanolamine-conjugated Atg8 localizes to the PAS. While the PAS is expanding, Atg8 is localized to the inner autophagosome membranes. Once the autophagosome maturation is complete, Atg8 is delivered to the vacuole along with the autophagosomes and recycled. While Atg8 is degraded, the GFP moiety is resistant to vacuolar proteases. Hence, the ratio of free GFP to GFP-Atg8 on a Western blot serves as a reporter of autophagy activity [Figure 1A]. To investigate whether autophagy induction is altered in the phosphatase mutants, we transformed WT, *ptc2Δ, ptc3Δ* and *ptc2Δ ptc3Δ* strains with a plasmid carrying GFP-Atg8 and induced autophagy by treating the cells with rapamycin to induce macroautophagy or with methyl methanesulfonate (MMS) to induce DNA damage and GTA (Eapen et al. 2017). We collected samples 4h after treatment, extracted proteins and immunoblotted them for GFP. The single mutants *ptc2Δ* and *ptc3Δ* showed WT levels of autophagy induction in response to either rapamycin or MMS, except for a small but statistically significant defect in *ptc3Δ* for GTA [Figure 1A]. In contrast, the *ptc2Δ ptc3Δ* double mutant was significantly defective for both rapamycin-induced autophagy and GTA [Figure 1B].

**Figure 1.**
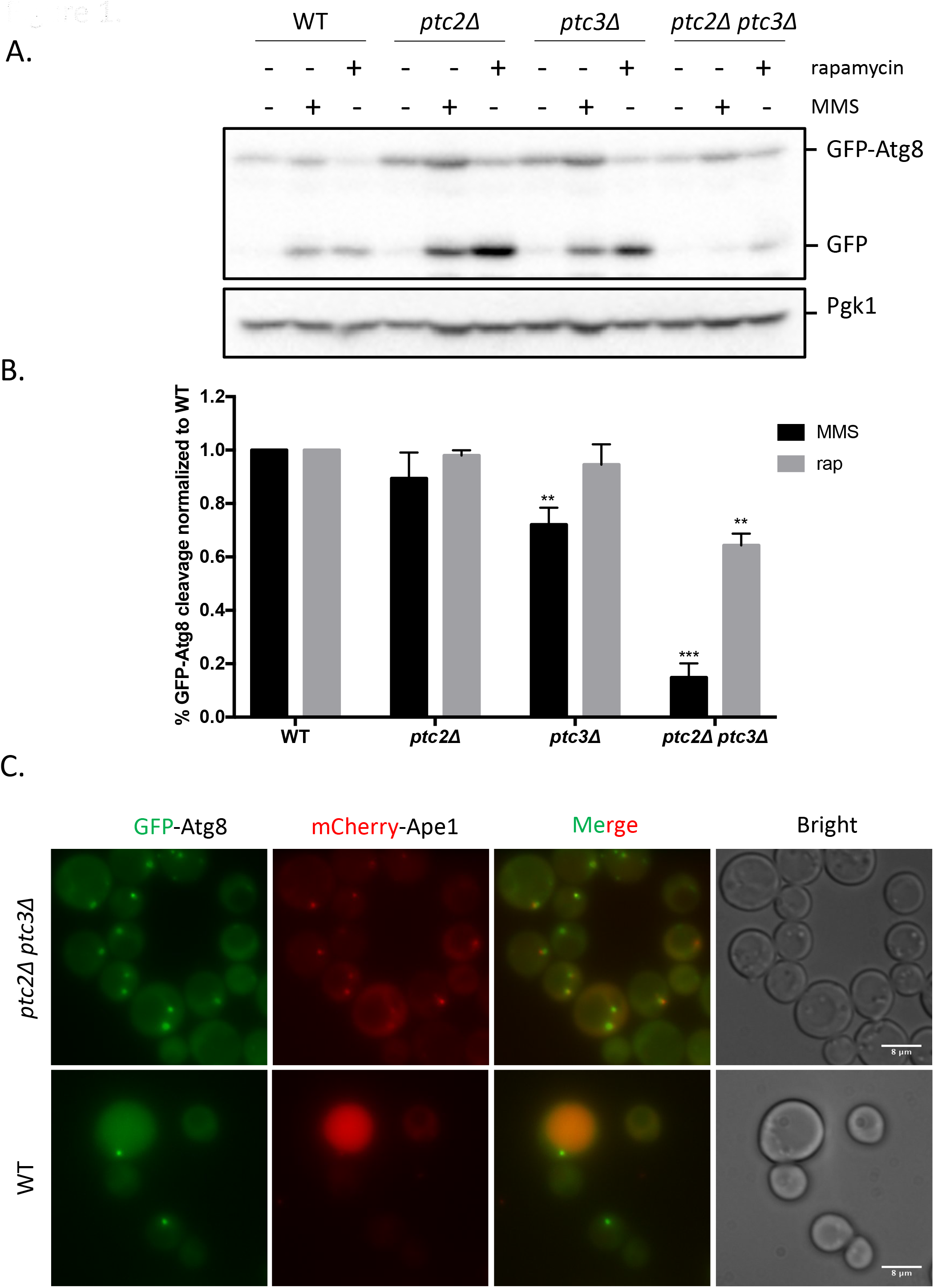
**A.** The effect of Ptc2 and Ptc3 on autophagy. WT and phosphatase mutants were grown in YEP-lac until they reached early exponential phase and treated with 200 ng/ml rapamycin or 0.04% MMS for 4h. Samples were blotted for GFP to measure the percentage of GFP-Atg8 processing, and for Pgk1 as a loading control. **B.** GFP-Atg8 cleavage was calculated as the ratio of free GFP band to the total GFP signal in the lane for 3 independent experiments, and then normalized to WT. Error bars represent standard error. **<0.002, ***<0.0002 **C.** The localization of GFP-Atg8 and mCherry-Ape1 in WT and phosphatase mutant strains after 1h of rapamycin treatment.

We confirmed these results by live cell fluorescent imaging of GFP-Atg8. In WT cells, after rapamycin treatment, GFP-Atg8 localized to the PAS and to the vacuole as previously shown (Kirisako et al. 1999). However, in the *ptc2Δ ptc3Δ* double mutant, the delivery of GFP-Atg8 to the vacuole was blocked significantly, and the vacuolar GFP signal in the double phosphatase mutant was significantly reduced compared to WT cells [Figure 1C]. We observed the same block after MMS treatment [Supplementary Figure 1]. GFP-Atg8 localization at the PAS after autophagy induction was more pronounced in the *ptc2Δ ptc3Δ* double mutant compared to WT. Moreover, in some cells, instead of one prominent GFP-Atg8 focus, there are multiple, less intense foci, that colocalize with mCherry-Ape1 foci 1h after rapamycin treatment [Figure 1C]. Taken together, these results suggest that Ptc2 and Ptc3 act redundantly to promote DNA damage-induced autophagy, as well as macroautophagy.

### PP2C-type phosphatases Ptc2 and Ptc3 promote cytoplasm-to-vacuole targeting pathway

After establishing that Ptc2 and Ptc3 promote macroautophagy, we asked if they are also involved in selective autophagy. The prototypic selective autophagy pathway in yeast is the cytoplasm-to-vacuole (Cvt) pathway (Lynch-Day and Klionsky 2010). To assay Cvt activity quantitatively, we used the Ape1 maturation assay (Torggler et al. 2017). Ape1 is synthesized in the cytoplasm as a precursor peptide (prApe1), and then delivered to the vacuole through the Cvt pathway for its maturation (mApe1) by the removal of its propeptide (Lynch-Day and Klionsky 2010). Maturation of Ape1 can be detected on a Western blot with an Ape1-specific antibody that recognizes both forms of the protein (Klionsky et al. 1992). We collected samples from WT and phosphatase null cultures after rapamycin and MMS treatment, as indicated before, and blotted them with anti-Ape1 antibody to assay Cvt activity. We found that deletion of *PTC2* and *PTC3* impaired Ape1 maturation under basal conditions, as well as after rapamycin or MMS treatments [Figure 2A]. We confirmed the inhibition of Ape1 delivery to the vacuole in the absence of Ptc2 and Ptc3 phosphatases by live cell microscopy, by using an N-terminus mCherry-tagged Ape1. We observed the accumulation of abnormally large mCherry-Ape1 foci at the PAS in the *ptc2Δ ptc3Δ* cells 4h after rapamycin treatment [Figure 2B]. Taken together, these results suggest that the delivery of Cvt vesicles in the absence of Ptc2 and Ptc3 is significantly impaired.

**Figure 2.**
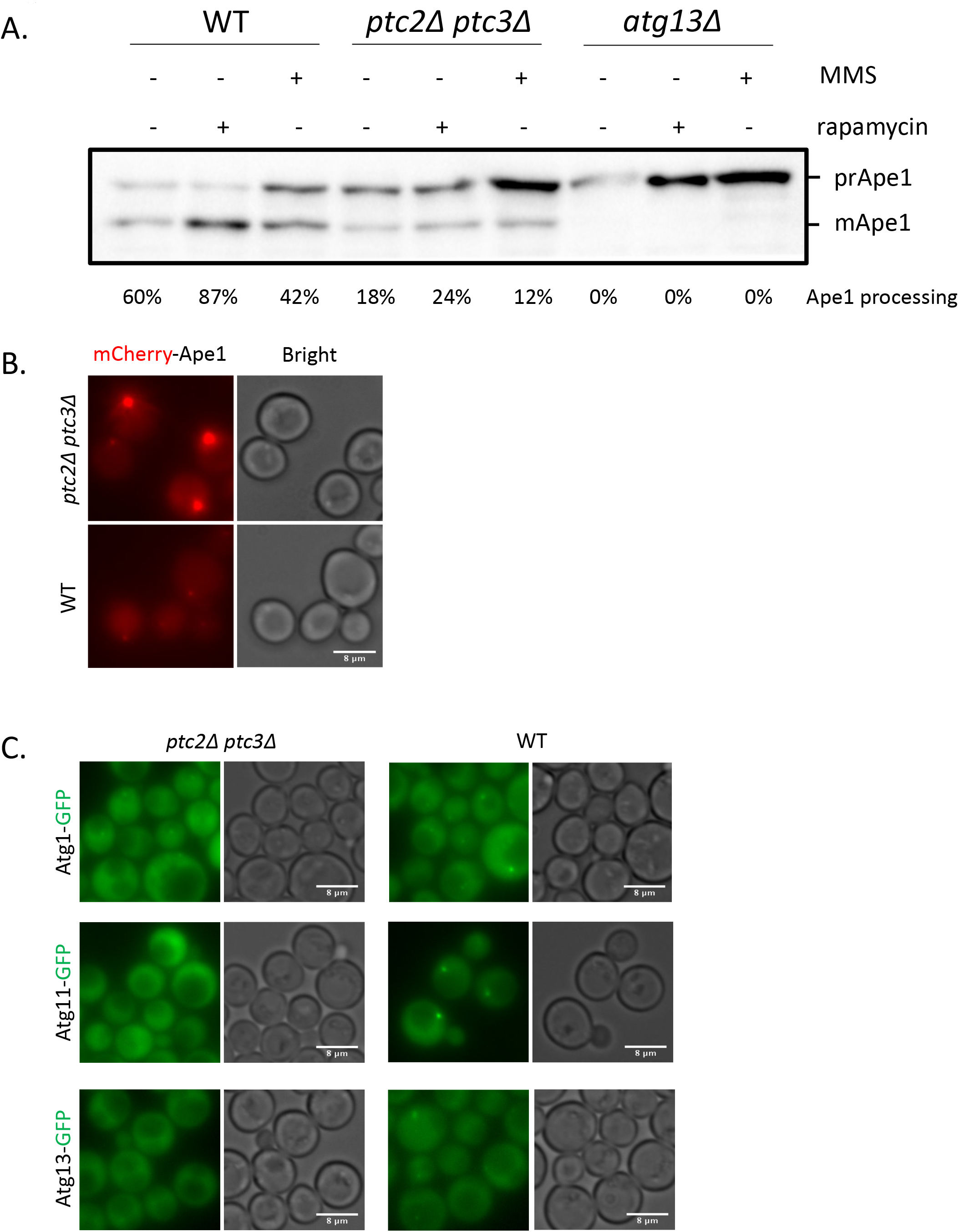
**A.** Ape1 processing in WT, *ptc2Δ ptc3Δ* and *atg13Δ* mutants after 4h rapamycin or MMS treatment. Ratio of mApe1 to total Ape1 signal for each lane is indicated as percent Ape1 processing below every lane. **B.** mCherry-Ape1 localization in *ptc2Δ ptc3Δ* and WT cells 4h after rapamycin treatment. **C.** The localization of Atg1-GFP, Atg11-GFP and Atg13-GFP in *ptc2Δ ptc3Δ* and WT cells growing in rich media.

The delivery of prApe1 to the vacuole is dependent on Atg1 and Atg13 (Kamada et al. 2000) and the adapter protein Atg11 (Yorimitsu and Klionsky 2005). To see if the Cvt defect of the *ptc2Δ ptc3Δ* cells is due to the mislocalization of these essential Cvt machinery, we assayed the localization of Atg1, Atg11 and Atg13 under nutrient replete conditions in WT and *ptc2Δ ptc3Δ* cells by using fluorescent live cell imaging. We observed weak Atg1 puncta in the absence of Ptc2 and Ptc3 phosphatases compared to WT. Moreover, in the absence of Ptc2 and Ptc3, Atg11 and Atg13 did not form visible puncta [Figure 2C], suggesting that phosphatase null cells are defective in recruitment of the ATG machinery required for proper Cvt function.

We also observed that the *ptc2Δ ptc3Δ* mutants have abnormal vacuoles by FM4-64M staining [Supplementary Figure 2], reminiscent of class-B *vps* mutants in their alveolar appearance (Vida and Emr 1995). Taken together, these data indicate that Ptc2 and Ptc3 are required for proper localization of autophagy proteins and the delivery of the Cvt vesicles to the vacuoles.

### Ptc2 and Ptc3 phosphatases regulate the localization of Atg proteins to the PAS

Atg1, Atg13 and Atg17 are the part of the core autophagy pathway; and are required for proper autophagy function (Kabeya et al. 2005, Kamada et al. 2000). Therefore, we asked if this core autophagy machinery is recruited to PAS properly in the absence of Ptc2 and Ptc3 phosphatases after autophagy induction. We used fluorescently-tagged Atg1, Atg11, Atg13 and Atg17 and assayed their localization with live cell fluorescent imaging 4h after rapamycin treatment. Similar to what we observed under nutrient-replete conditions, Atg1 was able to form weak puncta in the absence of Ptc2 and Ptc3 phosphatases [Figure 3A]. However, the number of cells that had Atg11, Atg13 and Atg17 foci were significantly reduced in the *ptc2Δ ptc3Δ* cells [Figure 3B, Supplementary Figure 3]. These results suggest that Ptc2 and Ptc3 are required for the proper recruitment of autophagy proteins to the PAS under nutrient starvation conditions.

**Figure 3.**
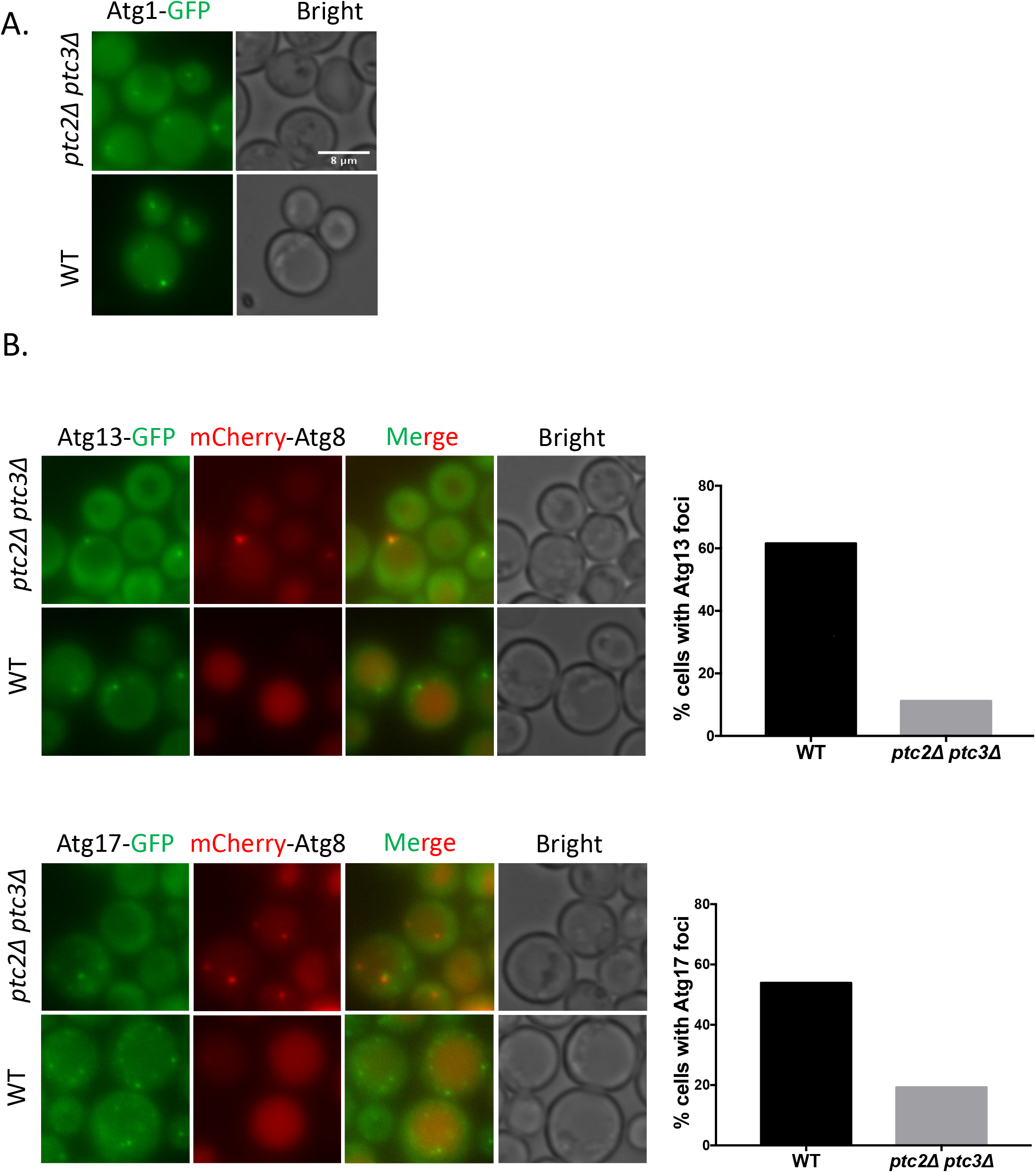
**A.** Atg1-GFP localization in *ptc2Δ ptc3Δ* and WT cells after 4h rapamycin addition. **B.** Atg13-GFP and Atg17-GFP localization in *ptc2Δ ptc3Δ* and WT cells 4h after rapamycin treatment. Percentage of cells with visible Atg13 or Atg17 foci were quantified and plotted. For each condition, more than 150 cells were counted.

### Atg1 is maintained in a hyperphosphorylated state in the absence of Ptc2 and Ptc3

Next, we investigated whether Ptc2 and Ptc3 act directly on TORC1 to regulate autophagy, since TORC1 signaling is inhibitory to macroautophagy. As a read-out of TORC1 activity, we assayed the phosphorylation levels of the AGC kinase Sch9, a direct target of TORC1 (Urban et al. 2007). We observe that both in WT and *ptc2Δ ptc3Δ* double mutant cells, Sch9 was phosphorylated under nutrient-replete conditions and was dephosphorylated upon rapamycin treatment [Supplementary Figure 4], indicating that the Ptc2 and Ptc3 phosphatases are likely to exert their effects on autophagy by acting on downstream targets of TORC1 rather than on TORC1 itself.

**Figure 4.**
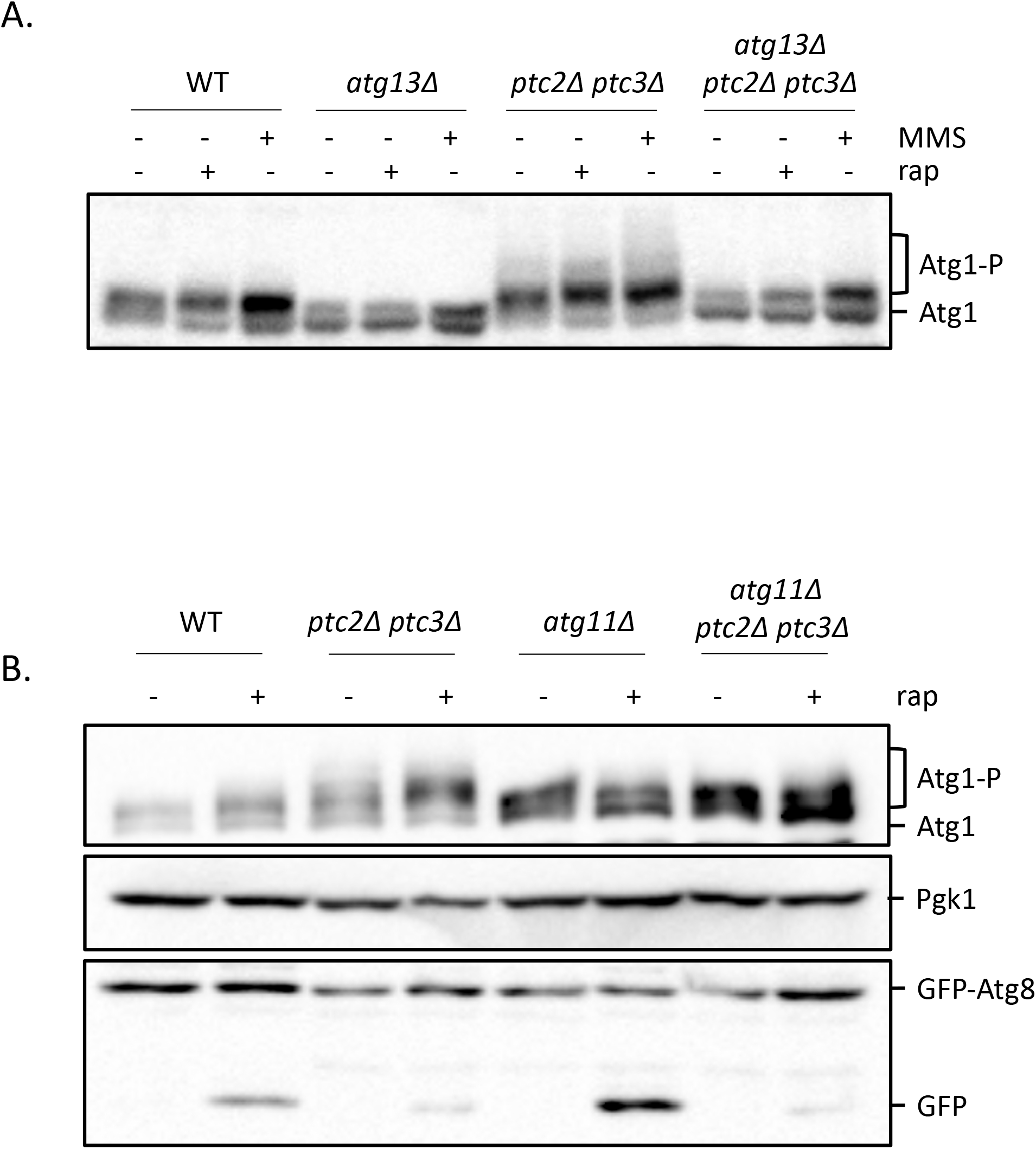
**A.** Atg1 hyperphosphorylation in WT, *atg13Δ, ptc2Δ ptc3Δ* and *atg13Δ ptc2Δ ptc3Δ* strains after 4h of rapamycin or MMS treatment. **B.** Atg1 hyperphosphorylation in WT, *ptc2Δ ptc3Δ, atg11Δ* and *atg11Δ ptc2Δ ptc3Δ* strains after 4 h of rapamycin treatment. Samples were blotted with anti-FLAG antibody for FLAG-Atg1 detection, Pgk1 for loading control and GFP to measure GFP-Atg8 cleavage.

Atg1 is regulated extensively by phosphorylations, some of which have been shown to be inhibitory for autophagic activity. Ptc2 and Ptc3 could promote autophagy by relieving these inhibitory phosphorylations. To monitor the phosphorylation of Atg1 in *ptc2Δ ptc3Δ* double mutant, we used a strain which carries an N-terminus 2XFLAG-tagged Atg1 allele (Kamber et al. 2015). Starvation-induced inhibition of TORC1 and PKA causes the activation of Atg1 through autophosphorylation (Kamada et al. 2000, Stephan et al. 2009). Atg1 autophosphorylation requires the presence of Atg13, as binding to dephosphorylated Atg13 stimulates Atg1’s kinase activity after autophagy induction (Yeh et al. 2010). We confirmed that Atg1 is phosphorylated 4h after rapamycin or MMS treatment, in an Atg13-dependent manner. In *ptc2Δ* and *ptc3Δ* single mutants, Atg1 was phosphorylated upon MMS or rapamycin treatment similar to WT [Supplementary Figure 5]. However, in the *ptc2Δ ptc3Δ* double mutant, we detected an accumulation of high molecular weight Atg1 bands in nutrient-replete conditions, as well as after the induction of autophagy. The appearance of these Atg1 bands in the double phosphatase mutant was Atg13-dependent, as the mobility of Atg1 in *atg13Δ ptc2Δ ptc3Δ* triple mutant was comparable to the *atg13Δ* single mutant [Figure 4A].

**Figure 5.**
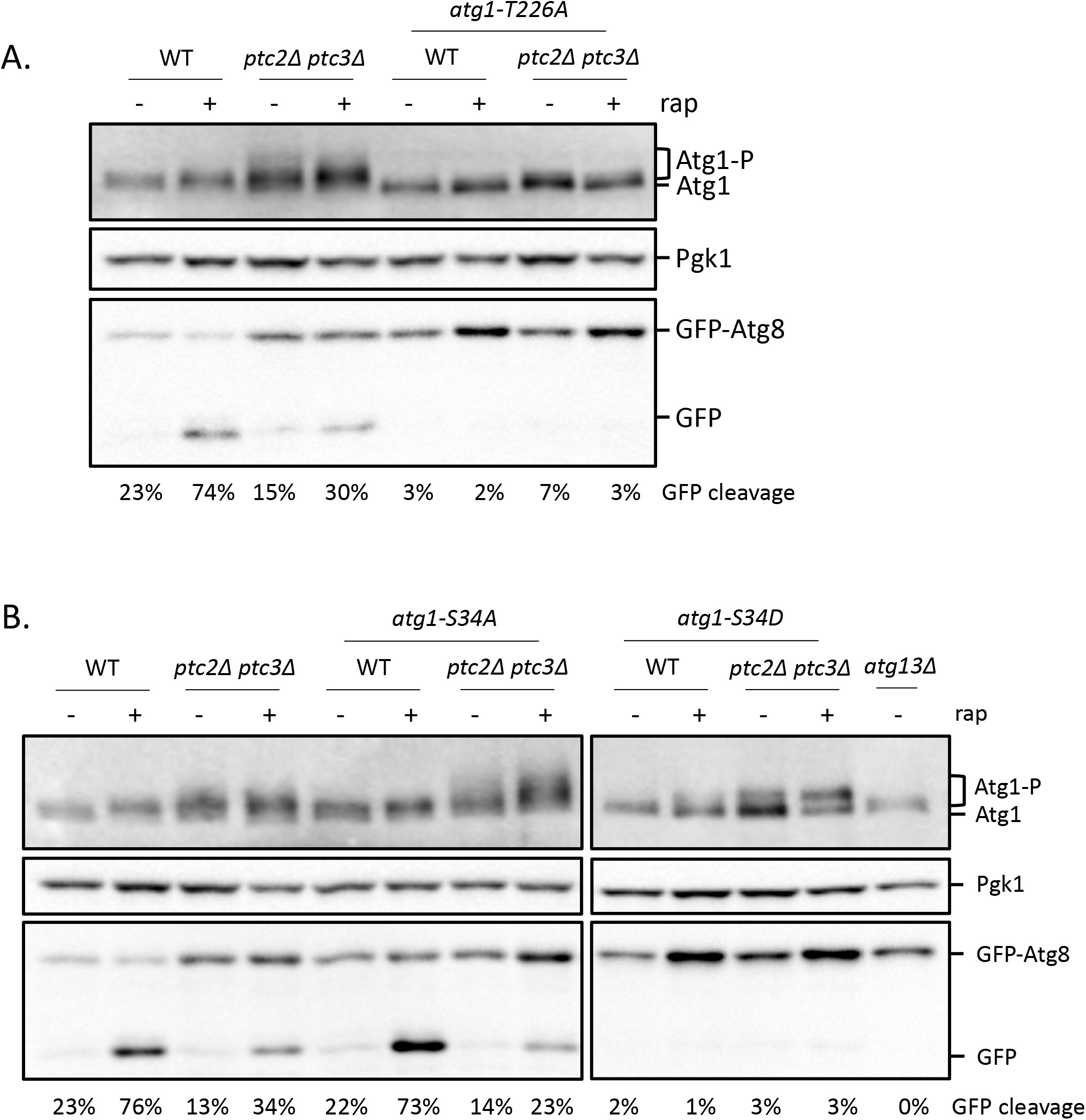
**A.** Atg1 hyperphosphorylation in phosphatase mutant strains that carry a *atg1-T226A* point mutation, which impairs Atg1’s kinase activity. Cells were treated with rapamycin for 4h, and samples collected were blotted with anti-FLAG to assay the phosphorylation status of Atg1, with anti-Pgk1 for loading control, and for GFP, to measure the GFP-Atg8 processing. Numbers below lanes indicate the GFP-Atg8 cleavage for the corresponding lane. **B.** Atg1 hyperphosphorylation in *atg1-S34A, atg1-S34D, atg1-S34D ptc2Δ ptc3Δ* and *atg1-S34D ptc2Δ ptc3Δ*.

Recent work from Denic lab showed that Atg1’s kinase activity under nutrient replete conditions is stimulated by binding to Atg19-prApe1 Cvt cargo complex, as well as the adapter protein Atg11 (Kamber et al. 2015). We asked if the hyperphosphorylated Atg1 species we observe in the absence of Ptc2 and Ptc3 phosphatases under nutrient replete conditions could be due to the overstimulation of Atg1 by Cvt cargo binding. To test this, we measured the phosphorylation levels of Atg1 in the *ptc2Δ ptc3Δ* cells lacking Atg11, Atg19 or Atg1 before or after rapamycin treatment. Interestingly, the appearance of hyperphosphorylated Atg1 species was dependent on Atg11 [Figure 4B], but not on Atg19 or Ape1 [Supplementary Figure 6]. Although in the *ptc2Δ ptc3Δ atg1lΔ* triple mutant, Atg1 hyperphosphorylation was reduced, this was not enough to rescue the macroautophagy defect of the *ptc2Δ ptc3Δ* as judged by the GFP-Atg8 processing [Figure 4B]. We conclude that Atg1 phosphorylation is counteracted by Ptc2 and Ptc3 phosphatases under nutrient-replete and starvation conditions.

**Figure 6.**
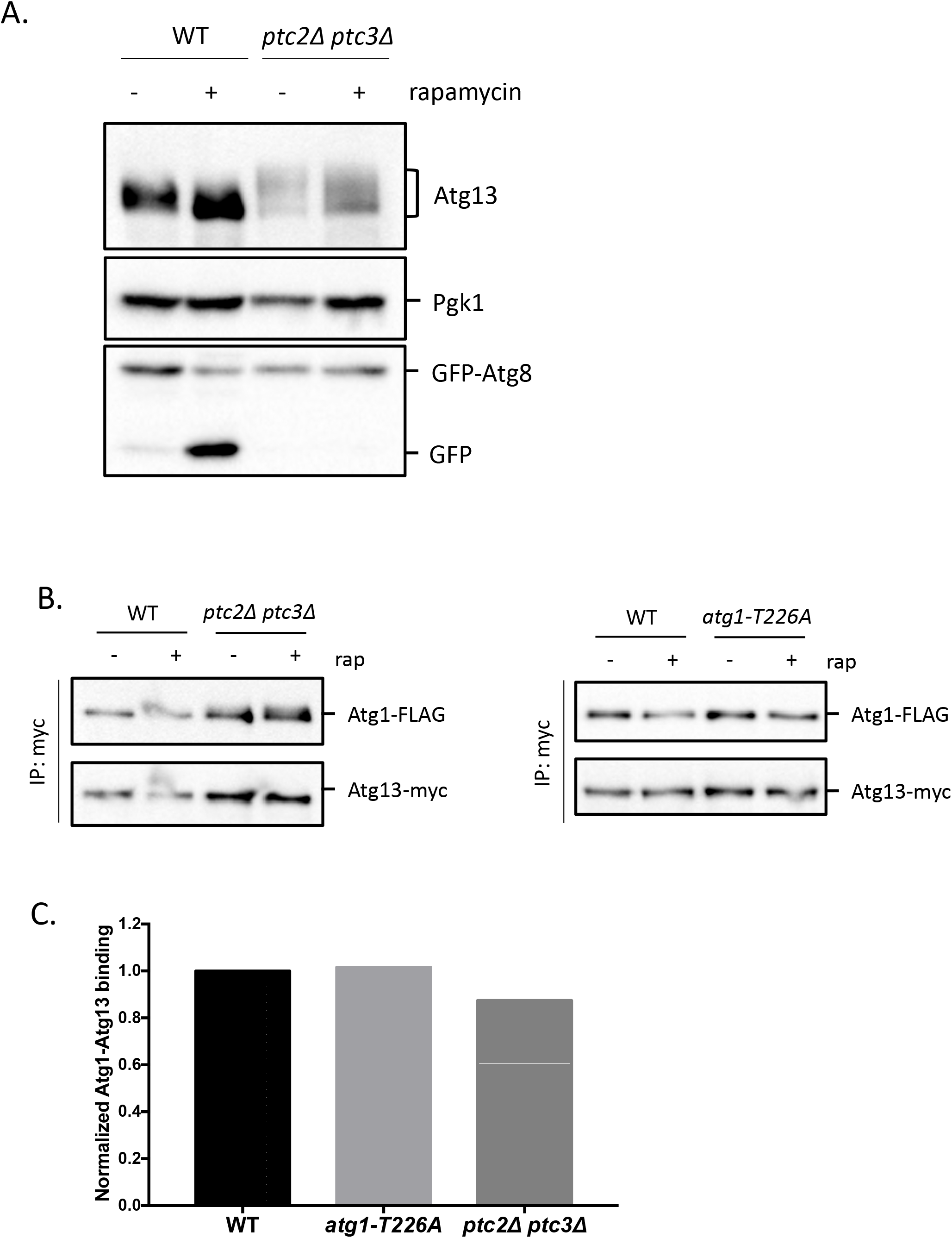
**A.** Atg1 and Atg13 hyperphosphorylation in WT and *ptc2Δ ptc3Δ* strains before and after 4h rapamycin treatment. Samples were collected and blotted for MYC-Atg13, Pgk1 and GFP. **B.** Atg1-Atg13 binding in *ptc2Δ ptc3Δ* and *atg1-T226A*. FLAG-Atg1 was coimmunoprecipitated with Atg13-MYC in WT, *ptc2Δ ptc3Δ* and *atg1-T226A* and blotted for FLAG, to detect Atg1 and for MYC, to detect Atg13. **C.** Normalized Atg1-Atg13 binding was calculated as a ratio of FLAG-Atg1 band intensity to Atg13-MYC band intensity at 0h for WT, *atg1-T226A* and *ptc2Δ ptc3Δ* and then normalized to WT.

Previous reports indicate that Atg1 autophosphorylation of residue T226 in the activation loop domain is required for Atg1’s kinase activity and proper autophagy induction (Kijanska et al. 2010, Yeh et al. 2010). An *atg1-T226A* point mutation blocks autophagy activity as well as the Atg1’s kinase activity (Yeh et al. 2010). We asked if the high mobility of Atg1 bands seen in *ptc2Δ ptc3Δ* double mutant were dependent on Atg1’s kinase activity. We created an *atg1-T226A* mutant at the endogenous locus by using CRISPR/Cas9 (see Methods) in WT and *ptc2Δ ptc3Δ* backgrounds that carry an epitope-tagged Atg1. As expected, the *atg1-T226A* point mutant abolished the autophagy-dependent autophosphorylation of Atg1 in WT. Additionally, the *atg1-T226A* substitution blocked the appearance of slower-migrating Atg1 bands in the *ptc2Δ ptc3Δ* double mutant [Figure 5A]. Autophagy was completely blocked in the strains carrying the *atg1-T226A* allele, in agreement with the previously shown role of this site in Atg1’s kinase activity (Yeh et al. 2010). These data demonstrate that Atg1 is extensively autophosphorylated in the absence of PP2C type phosphatases Ptc2 and Ptc3 irrespective of autophagy induction, and PP2C-type phosphatases counteract these phosphorylations.

In addition to T226, Atg1 is regulated by phosphorylation of serine 34; however, the phosphorylation of Atg1-S34 is inhibitory for autophagy as well as for Atg1’s kinase activity (Yeh et al. 2011a). To see if Ptc2 and Ptc3 exert their effect on Atg1 by reversing Atg1-S34 phosphorylation, we created endogenous *atg1-S34A* and *atg1-S34D* point mutants by using CRISPR/Cas9 in WT and phosphatase null backgrounds. We induced autophagy by the addition of rapamycin for 4h and measured the GFP-Atg8 processing together with Atg1 mobility shift. Our data show that autophagy is impaired in *atg1-S34D* point mutant as previously shown (Abeliovich et al. 2000), but this substitution is not sufficient to fully eliminate the autophosphorylation of Atg1 after autophagy induction [Figure 5B]. The *atg1-S34D* substitution reduced the levels of hyperphosphorylated Atg1 detected in *ptc2Δ ptc3Δ*, which could be due to the decreased level of Atg1 kinase activity observed in the S34D mutant (Yeh et al. 2011a), and hence, the defects in Atg1 autophosphorylation observed in *ptc2Δ ptc3Δ* double mutants. In contrast, the autophagy-proficient *atg1-S34A* point mutant failed to rescue the autophagy defect in the double phosphatase mutants, suggesting that additional Atg1 phosphorylation sites are involved in this regulation or that S34 specifically is not regulated by PP2C phosphatases.

Prior research suggested that the more slowly-migrating Atg1 species observed post-autophagy induction is due to the phosphorylation of site S390 and that the Atg1 phospho-shift detected upon autophagy induction is lost in an *atg1-S390A* point mutant (Yeh et al. 2011a). We asked if the phosphorylation of S390 contributed to the higher migrating Atg1 bands in the *ptc2Δ ptc3Δ* double mutant. We created *atg1-S390A* and *atg1-S34A* single and double mutants with CRISPR/Cas9 in WT and *ptc2Δ ptc3Δ* strains and assayed the phosphorylation status of Atg1 before or after the induction of autophagy by rapamycin treatment. We observed a more prominent rapidly-migrating band in the *atg1-S34A,S390A ptc2Δ ptc3Δ* quadruple mutant; however, neither the *atg1-S34A* and *atg1-S390A* single mutants, nor the *atg1-S34A,S390A* double mutant was able to rescue the autophagy defect of *ptc2Δ ptc3Δ* [Supplementary Figure 7]. Collectively, these results suggest that Ptc2 and Ptc3 are required for Atg1 dephosphorylation; however, the S34 and S390 sites that were previously implicated in Atg1 regulation are either not directly targeted by Ptc2 and Ptc3, or there are additional sites, yet to be identified, which play a role in Atg1 regulation.

**Figure 7.**
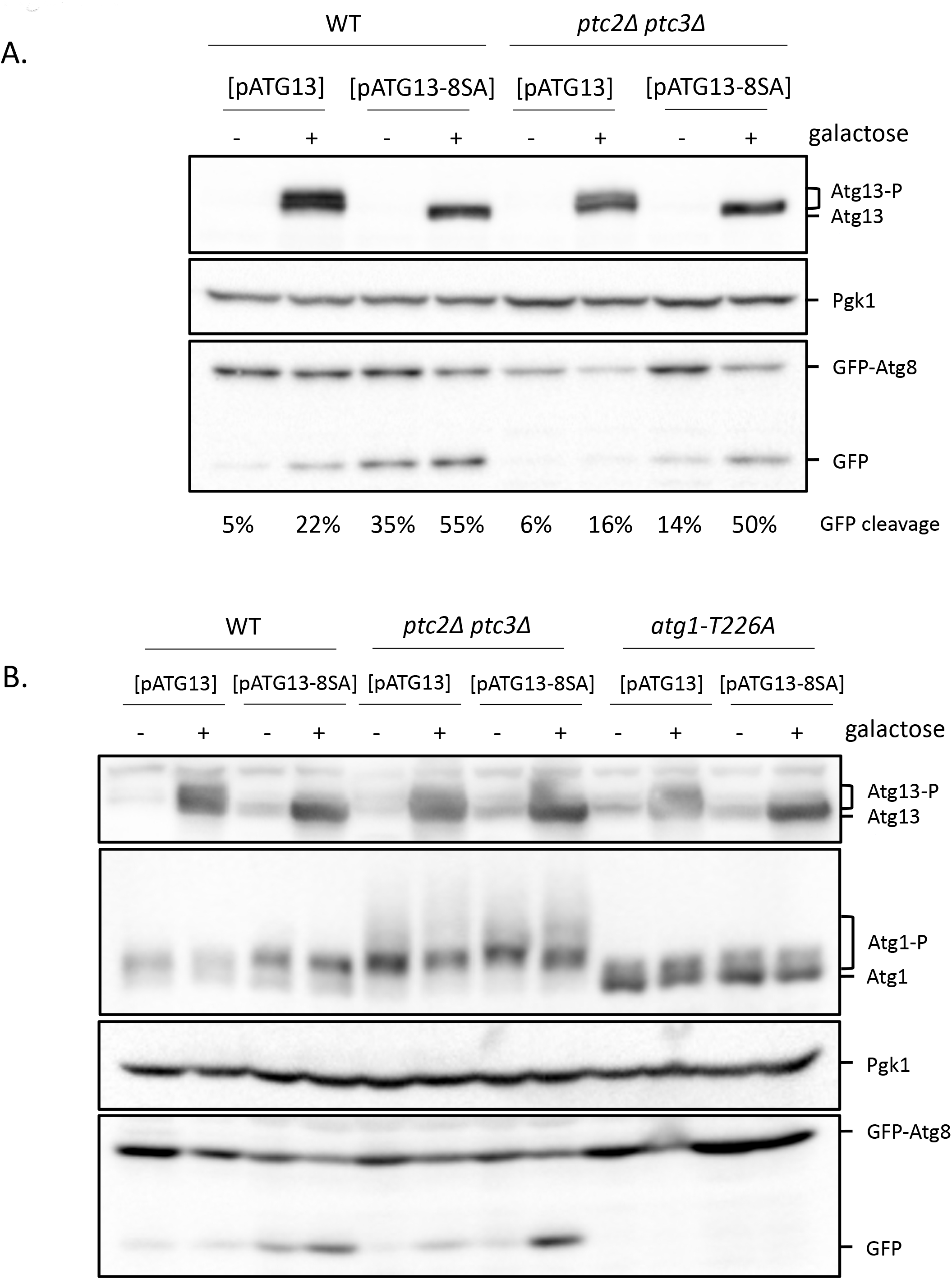
**A.** Overexpression of *ATGl3-8SA* in WT and *ptc2Δ ptc3Δ*. Strains containing WT *ATGl3* or *ATGl3-8SA* centromeric plasmids under galactose-inducible promoters were grown and induced with 2% galactose for 4h. Samples were collected and blotted with anti-Atg13 antibody, anti-Pgk1 antibody and anti-GFP antibody. The percent GFP-Atg8 processing for every lane is indicated below the corresponding lane. **B.** Atg1 hyperphosphorylation and GFP-Atg8 processing after the overexpression of WT *ATGl3* or *ATGl3-8SA* in WT, *ptc2Δ ptc3Δ* and *atg1-T226A* strains. FLAG-Atg1 strains carrying the WT *ATGl3* or *ATGl3-8SA* galactose-inducible plasmids were grown and induced as explained previously. Samples were collected and blotted for Atg13, FLAG (for Atg1 detection) and for GFP.

### Atg13 is hyperphosphorylated in *ptc2Δ ptc3Δ* double mutant

As mentioned previously, Atg13 is hyperphosphorylated prior to autophagy induction by the action of TORC1 and PKA kinases. Upon nutrient starvation or rapamycin treatment, TORC1 activity is downregulated and Atg13 is rapidly dephosphorylated. This step is required for proper autophagy induction, since overexpression of a constitutively active *ATG13-8SA* allele is sufficient to induce autophagy in cells carrying a wild-type *ATG13* gene (Kamada et al. 2010). We hypothesized that one way by which Ptc2 and Ptc3 could promote autophagy is by mediating the dephosphorylation of Atg13 during initiation of autophagy. If that were the case, we would expect to detect constitutively phosphorylated Atg13 in *ptc2Δ ptc3Δ* mutants, regardless of the status of TORC1. To test this hypothesis, we aimed to detect the Atg13 phosphorylation in the phosphatase mutants. To be able to detect the endogenous Atg13, we created a strain that bears a genomic 10XMYC-Atg13. We assayed the phosphorylation of Atg13 in *ptc2Δ ptc3Δ* double mutant cells prior to autophagy induction and after 4h rapamycin treatment by immunoblotting. We detected an accumulation of apparently more hyperphosphorylated Atg13 prior to autophagy induction in the *ptc2Δ ptc3Δ* double mutant compared to WT [Figure 6A]. After autophagy induction, Atg13 was only partially dephosphorylated in *ptc2Δ ptc3Δ* cells, whereas in WT cells Atg13 was fully dephosphorylated upon autophagy induction as expected. Autophagy induction in the *ptc2Δptc3Δ* double mutant cells that contain an epitope-tagged Atg13 allele was blocked as expected [Figure 1A, Figure 6A]. These results suggest that the phosphatases Ptc2 and Ptc3 are involved in dephosphorylation of Atg13. We also observed that Atg13 was dephosphorylated in the *atg1-T226A* strain just like WT, suggesting that the dephosphorylation of Atg13 is independent of autophagy flux and Atg1’s kinase activity [Supplementary Figure 8].

**Figure 8.**
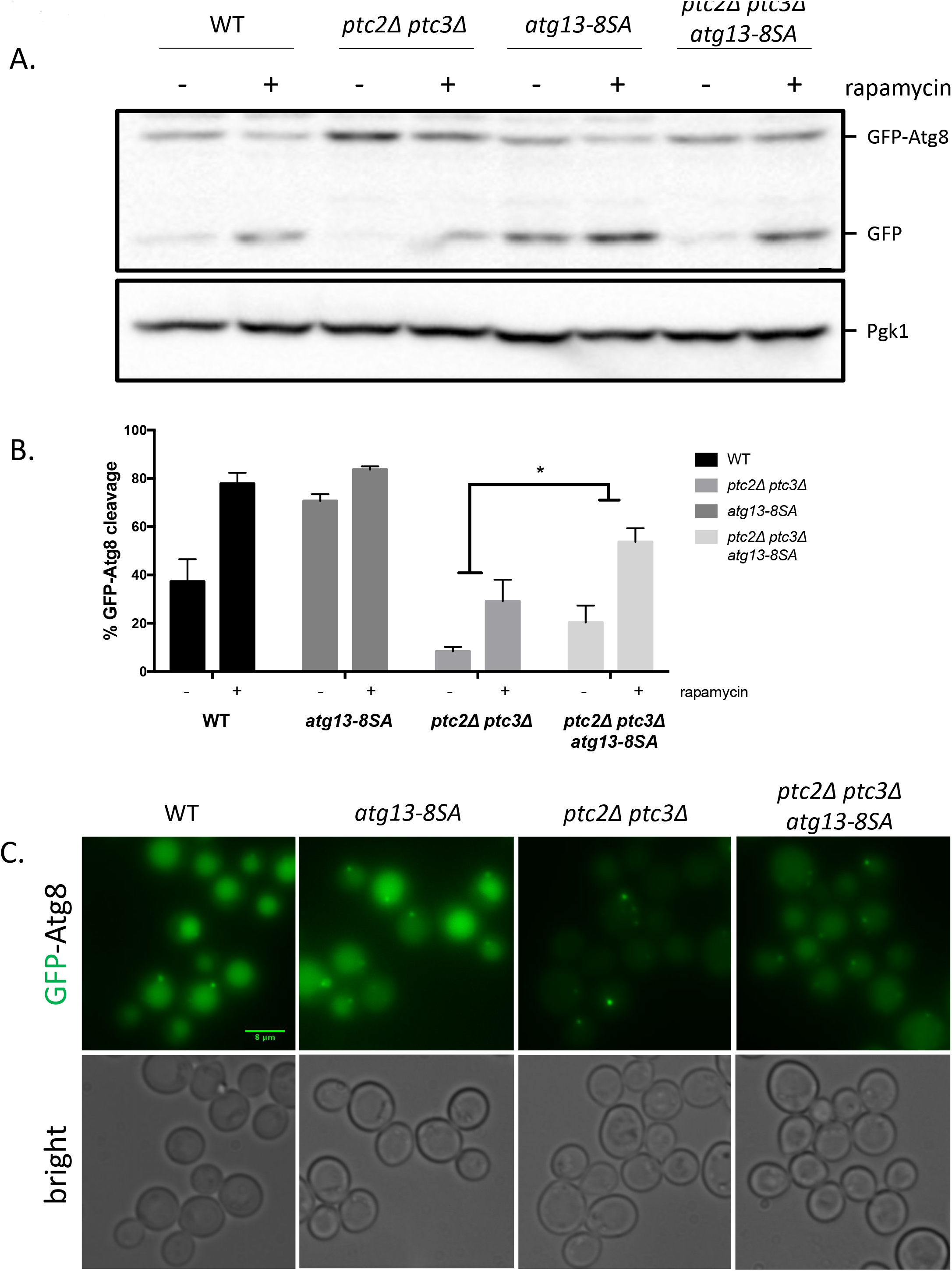
**A.** GFP-Atg8 processing in strains that carry *atg13-8SA* at endogenous locus after 4h rapamycin treatment. Samples were collected and blotted as described in Figure 1. **B.** Quantification of GFP-Atg8 processing in WT, *ptc2Δ ptc3Δ, atg13-8SA* and *ptc2Δ ptc3Δ atg13-8SA* strains. GFP-Atg8 processing in these strains were calculated from 3 independent experiments, averaged and plotted. Error bars represent standard error. P = 0.04. C. GFP-Atg8 localization in WT, *ptc2Δ ptc3Δ, atg13-8SA* and *ptc2Δ ptc3Δ atg13-8SA* strains 4h after rapamycin treatment.

Atg1 and Atg13 interact prior to autophagy induction (Kraft et al. 2012); however, hyperphosphorylated Atg13 has been shown to have weaker affinity for Atg1 (Kabeya et al. 2005, Kamada et al. 2000). Moreover, the binding of Atg13 stimulates Atg1’s kinase activity. One explanation for the autophagy defect of the *ptc2Δ ptc3Δ* cells is the inability to form a functional Atg1 complex due to the accumulation of hyperphosphorylated Atg1 and Atg13 species. To test this, we co-immunoprecipitated Atg13-13XMYC together with 2XFLAG-Atg1 [Figure 6B]. We found comparable levels of association of Atg1 with Atg13 in WT and phosphatase null strains as well as in a strain that carries the kinase defective *atg1-T226A* prior to autophagy induction [Figure 6C]. These results demonstrate that the hyperphosphorylated Atg1 and Atg13 species observed in the phosphatase null strain still can form a complex when *TORC1* is active, and that the autophagy defect seen in the *ptc2Δ ptc3Δ* null strain is not due to a defect in Atg1-Atg13 complex formation.

### The autophagy defect of the *ptc2Δ ptc3Δ* null cells is rescued by expression of constitutively active Atg13

As mentioned previously, the overexpression of the constitutively active *ATG13-8SA* is sufficient to induce autophagy that is uncoupled from TORC1 activity (Kamada et al. 2010). We asked if this would also be sufficient to bypass the autophagy defect of *ptc2Δ ptc3Δ* double mutant. Our data show that the overexpression of the constitutively active *ATG13-8SA* allele in a PP2C phosphatase null strain promoted autophagy to near-WT levels [Figure 7A]. Based on these data, we argue that the defect seen in the phosphatase null strain is directly due to a failure to dephosphorylate Atg13.

Overexpression of the constitutively active *ATG13-8SA* failed to reduce the hyperphosphorylation of Atg1 in *ptc2Δ ptc3Δ* [Figure 7B]. Moreover, the overexpression of constitutively active *ATG13-8SA* allele in the *atg1-T226A* background did not restore the autophagy defect of *atg1-T226A*, as opposed to the rescue observed in the *ptc2Δ ptc3Δ* strain, as we and others have previously shown (Kamada et al. 2010). Taken together, these results suggest that despite the extensive autophosphorylation, the kinase activity and Atg13 binding affinity of Atg1 are not impaired in the absence of Ptc2 and Ptc3. These data demonstrate that the impaired autophagy activity in the *ptc2Δ ptc3Δ* mutant strain is due to a defect in the dephosphorylation of Atg13 and not due to an inherent defect in Atg1’s kinase activity.

We noticed that overexpression of WT *ATG13* or presence of *ATG13-8SA* without the overexpression resulted in an increase in autophagy induction [Figure 7A and B]. The overexpression of *ATG13* has been shown to exacerbate the activating autophosphorylation of Atg1 at site T226A (Yeh et al. 2010), suggesting that Atg13 abundance can also regulate autophagy. To distinguish between these possibilities, we integrated the *atg13-8SA* allele into the genome, under the control of Atg13’s endogenous promoter in WT and *ptc2Δ ptc3Δ* backgrounds. As expected, the *atg13-8SA* strain showed increased levels of GFP-Atg8 cleavage even prior to rapamycin treatment [Figure 8A]. The levels of autophagy flux increased only modestly upon rapamycin treatment in this strain, once again reinforcing the idea that the phosphorylation of Atg13 at these 8 sites by the action of TORC1 kinase is the main inhibitory control over autophagy induction. The GFP-Atg8 cleavage observed in the *ptc2Δ ptc3Δ atg13-8SA* mutant was significantly higher compared to *ptc2Δ ptc3Δ;* however, introducing the *atg13-8SA* mutations did not restore WT levels of autophagy to the phosphatase null strain [Figure 8B]. Moreover, we observed an intermediate level of GFP localization to the vacuole in the *ptc2Δ ptc3Δ atg13-8SA* triple mutant, agreeing with the GFP-Atg8 cleavage assay results [Figure 8C]. These data suggest that Ptc2 and Ptc3 are involved in the regulation of other proteins in the autophagy pathway in addition to Atg13.

## DISCUSSION

We have previously shown that DNA damage induces a targeted autophagy pathway, termed GTA. This pathway is selective, and requires all the core ATG machinery, in addition to the scaffold protein Atg11 (Eapen et al. 2017). Moreover, the magnitude of GTA is positively correlated with the magnitude of DNA damage. The PP2C phosphatases Ptc2 and Ptc3 were previously shown to be negative regulators of DNA damage checkpoint response, specifically in regulating the autophosphorylation of Rad53/CHK2 (Guillemain et al. 2007, Leroy et al. 2003). Thus, in the absence of Ptc2 and Ptc3, even after cells have repaired DNA damage, they fail to exit from the checkpoint arrest and deactivate Rad53. Rad53 activation has also been shown to be critical for the induction of GTA (Eapen et al. 2017). For this reason, we expected to observe an increase in the GTA response in the absence of these phosphatases. However, we found a significant decrease in rapamycin-induced macroautophagy, as well as in GTA suggesting that Ptc2 and Ptc3 redundantly *promote* autophagy. In addition to the macroautophagy defect, *ptc2Δ ptc3Δ* mutants exhibit a strong Cvt defect. Ape1 maturation observed under nutrient-replete conditions is almost fully blocked in the absence of Ptc2 and Ptc3. We also observe very large Ape1 complexes after rapamycin treatment, and a defect in the recruitment of proteins that are essential to Cvt function *ptc2Δ ptc3Δ* mutant strain. Given that Ptc2 and Ptc3 do not act directly on TORC1, based on our data, we conclude that Ptc2 and Ptc3 modulate the core autophagy machinery downstream of TORC1.

To identify the targets of Ptc2 and Ptc3 among the core ATG machinery, we turned our attention to the Atg1 kinase complex. We observe that Atg1 is maintained in an hyperphosphorylated state in the *ptc2Δ ptc3Δ* double mutant. The appearance of the hyperphosphorylated Atg1 species is Atg13 and Atg11-dependent. Moreover, reducing the kinase activity of Atg1 by introducing a T226A substitution blocked the appearance of these bands. Based on these results, we suggest that the basal level binding between the Atg1, Atg13 and Atg11 under nutrient-replete conditions stimulates the kinase activity of Atg1 as previously observed (Kamber et al. 2015), and this results in extensive autophosphorylation of Atg1. Under normal circumstances, these phosphorylations are counteracted by Ptc2 and Ptc3, but in the absence of these phosphatases, Atg1 is maintained in a hyperphosphorylated state. Interestingly, the aberrant hyperphosphorylated Atg1 species in *ptc2Δ ptc3Δ* mutants are not dependent on Atg19-prApe1 cargo complex. We do not rule out that under normal conditions, Atg1’s kinase activity is stimulated by these proteins; although, we also observed that in the absence of Ptc2 and Ptc3, Atg11 is not recruited to the PAS. This implies that in the absence of Ptc2 and Ptc3 phosphatases, Atg1-Atg19-prApe1 binding is impaired due to the failure to recruit Atg11. Previous research showed that in the absence of Atg11, Atg1 and Atg13 formed a complex without Atg19, supporting this hypothesis (Kamber et al. 2015). Our data also imply that Atg1-Atg13 and Atg11 can interact without being localized to the PAS, since the localization of Atg13 and Atg11 is significantly impaired in the absence of Ptc2 and Ptc3 phosphatases.

PAS organization is a highly regulated process. Previous work demonstrated that the localization of the scaffold protein Atg17 to PAS is a prerequisite for Atg13 and Atg1 recruitment (Suzuki et al. 2007). In agreement, our data show that Atg1, Atg13 and Atg17 localization to PAS in the absence of Ptc2 and Ptc3 mutants is significantly impaired. This defect could be due to the regulation of Atg17 directly by Ptc2 and Ptc3 phosphatases, or a defect in Atg13-Atg17 binding. Indeed, Atg13-Atg17 binding is improved upon autophagy induction and Atg13 dephosphorylation (Kabeya et al. 2005). Therefore, it is possible that Atg13-Atg17 binding is weaker in the absence of Ptc2 and Ptc3 phosphatases, due to the hyperphosphorylation of Atg13.

After establishing that Atg1 is extensively autophosphorylated in *ptc2Δ ptc3Δ* mutant strain, we focused on previously characterized Atg1 phosphorylation sites to identify the sites that are involved in the regulation of Atg1 by these phosphatases. The phosphorylation of site Atg1-S34 has previously been characterized as inhibitory to autophagy activity (Yeh et al. 2011a). If the autophagy defect seen in the phosphatase null strain is due to the persistence of the inhibitory phosphorylation of Atg1 at site S34, introducing an alanine substitution should rescue this defect. However, *ptc2Δ ptc3Δ atg1-S34A* triple mutant is as defective as the *ptc2Δ ptc3Δ* double mutant for GFP-Atg8 processing, suggesting that the autophagy defect seen in the phosphatase mutants is not only due to the inhibitory phosphorylation of S34; additional sites are involved in the regulation. The *atg1-S34D* substitution, as well as the *atg1-S34A,S390A* substitutions decreased the abundance of the higher molecular weight Atg1 bands detected in the phosphatase null strain. But this could be due to the decrease in Atg1’s kinase activity in these mutants, hence, decreased levels of Atg1 autophosphorylation (Yeh et al. 2011a). Alternatively, the effect of Ptc2 and Ptc3 on Atg1 could be indirect.

Atg13, an essential component of the core autophagy machinery, is kept in a hyperphosphorylated and inhibited state under nutrient-replete conditions by TORC1 and PKA-dependent phosphorylations, and its dephosphorylation is critical for the autophagy induction. The phosphatases that are responsible for regulating Atg13 remain unidentified. Our data show that, in the absence of PP2C phosphatases, Atg13 is hyperphosphorylated irrespective of the TORC1 status, indicating that Ptc2 and Ptc3 are involved in dephosphorylation of Atg13. Previous work showed that the overexpression of constitutively active *ATG13-8SA* in a cell carrying a chromosomal *ATG13* allele can cause autophagy induction when TORC1 is active, in an Atg1-dependent manner (Kamada et al. 2010). The overexpression of this allele was also able to induce autophagy in the *ptc2Δ ptc3Δ* strain. Here we show the overexpression of *ATG13-8SA* is also able to sufficient to induce autophagy in the *ptc2Δ ptc3Δ* strain. Moreover, a single integrated *atg13-8SA* allele is able to partially rescue the autophagy defect of the phosphatase null strain. The *atg13-8SA* mutant induces autophagy under nutrient-replete conditions whereas the *ptc2Δ ptc3Δ atg13-8SA* strain does not. These results indicate that the autophagy defect observed in the absence of Ptc2 and Ptc3 is mainly due to a defect in the dephosphorylation of Atg13, but Atg13 is not the sole target of these phosphatases. A mass spectrometry study by Fujioka et al. suggested that there are 51 phospho-sites on Atg13 whose phosphorylation levels decrease after rapamycin treatment (Fujioka et al. 2014, Kamada et al. 2010). It is also possible that Ptc2 and Ptc3 regulate additional sites on Atg13 in addition to the sites that are targeted by TORC1.

Atg13 and the kinase activity of Atg1 are essential for Cvt function; however, Cvt can occur under nutrient rich conditions without TORC1 inhibition or without Atg13 dephosphorylation (Kamada et al. 2000, Yeh et al. 2010). Therefore, the Cvt defect of the *ptc2Δ ptc3Δ* double mutant is unlikely to be dependent on the accumulation of hyperphosphorylated Atg13. We show that in the absence of Ptc2 and Ptc3 phosphatases, the localization of Atg1, Atg11 and Atg13 are impaired, all of which are required for Cvt function, suggesting that there is a fundamental defect in PAS organization. It is possible that Ptc2 and Ptc3 modulate proteins that play an important role for selective autophagy but dispensable for the starvation-induced autophagy, such as Atg11 or Atg19 (S. V. Scott et al. 2001, Yorimitsu and Klionsky 2005).

We conclude that the Ptc2 and Ptc3 phosphatases are positive regulators of autophagy. They are involved in the dephosphorylation of Atg13 to stimulate the kinase activity of the Atg1 kinase complex to allow proper autophagy induction. The autophagy defect of the phosphatase null strain can be rescued by the overexpression of constitutively active *ATG13-8SA* allele or expressing the *atg13-8SA* under its endogenous promoter. Moreover, according to our results, Atg1 is constitutively autophosphorylated under nutrient replete conditions. These phosphorylations are not inhibitory to its kinase activity and are counteracted by PP2C type phosphatases Ptc2 and Ptc3. The effect of these autophosphorylations on Atg1 regulation is still not known.

## MATERIALS AND METHODS

### Yeast media, strains and plasmids

*Saccharomyces cerevisiae* strains used in this study are derivatives of JKM179 or BY4741 (**Supplementary Table 3**). Plasmids used in the study are listed in **Supplementary Table 1**. Deletions of ORFs were created by using the one step PCR homology cassette amplification and transformed using high efficiency yeast transformation methods (Amberg et al. 2006, Wach et al. 1994). The primer sequences used for the deletions are available upon request. Atg1 point mutants were created by using CRISPR/Cas9 as previously explained (Anand et al. 2017). The sequences of the single-stranded oligonucleotide DNA (ssODN) templates used to create CRISPR/Cas9-mediated point mutants are indicated in **Supplementary Table 2**, the CRISPR/Cas9 plasmids are indicated in **Supplementary Table 1**. 2XFLAG-Atg1 in the Atg13-13xMYC strain was created by targeting the N-terminus of Atg1 with CRISPR/Cas9 and using a PCR-amplified fragment as template. N-terminus mCherry-tagged Ape1 and N-terminus 10xMYC-tagged Atg13 were created by targeting the N-terminus of the ORFs with CRISPR/Cas9. The Atg13-13xMYC epitope-tagged strain was created by transforming cells with PCR-amplified cassettes that contain homology to Atg13 C-terminus (Longtine et al. 1998). The GFP-Atg8, *GAL-ATG13, GAL-ATG13-8SA*, mCherry-Ape1 and 2GFP plasmids were a gift from Yoshiaki Kamada, National Institute of Basic Biology, Okazaki, Japan (Kamada et al. 2010). The 2XFLAG-Atg1 VDY630 strain was a gift from Vladimir Denic, Harvard University, Cambridge, MA, USA (Kamber et al. 2015). C-terminus 2XGFP tagging of the Atg1, Atg11, Atg13 and Atg17 was performed as previously described (Li et al. 2015). Yeast strains were grown in YEPD (1% yeast extract, 2% peptone, 2% dextrose) for normal growth conditions.

### GFP-Atg8 processing assay

Cells were grown in YEPD overnight, washed into YEP-lactate (YEP containing 3% lactic acid) and inoculated into YEP-lac so that they reach exponential phase. For galactose induction experiments, cells were treated with a final concentration of 2% galactose. We used MMS at a final concentration of 0.04% to induce GTA, or rapamycin at a final concentration of 200ng/ml to induce macroautophagy.

### Sample collection and Western blotting

Cells were harvested from 50 ml of the early exponential phase culture (1-10 x 10^6^ cells/ml) by centrifugation. Cell pellets were washed in 1 ml of 20% TCA and quick-frozen on dry ice. TCA protein extraction was carried out as previously described (Eapen et al. 2017). For pulldown experiments, approximately 500 μg of cell pellet was harvested and washed with ice cold 1 ml 50 mM Tris, 5 mM EDTA solution. The samples were kept on dry ice. For protein extraction, cells were resuspended in lysis buffer (50 mM HEPES pH7.5, 150 mM NaCl, 2 mM EDTA, 0.5% IGEPAL), supplemented with protease inhibitors. Cell lysis was performed by vortexing with acid-washed glass beads. The crude extract was collected and cleared by centrifugation at maximum speed twice for 15 min at 4^0^C. An equal volume of the cleared lysate was mixed with Protein G-agarose beads (Roche) pre-incubated with anti-MYC antibodies. After the overnight incubation, the beads were washed twice with 1ml lysis buffer containing protease inhibitors, and the samples were eluted in 100 μl 6X Laemmli buffer containing 0.9% B-mercaptoethanol by boiling at 95^0^C for 5 min. Samples were run on 6% gels for the detection of hyperphosphorylated Atg1 and Atg13 species, 7% gels for Sch9 and Ape1 detection, and 10% gels for GFP-Atg8, Pgk1, and Atg13 overexpression assays. Blotting was performed by using anti-GFP antibody (AbCam ab-6556), anti-FLAG antibody (Millipore Sigma A2220), anti-Pgk1 antibody (AbCam ab113687), anti-MYC antibody (AbCam 9E10), anti-Atg13 antibody, anti-Ape1 antibody, and anti-Sch9 antibody. Anti-Atg13 and anti-Ape1 antibodies were a gift from Dr. Daniel Klionsky (Klionsky et al. 1992, Miller-Fleming et al. 2014). Anti-Sch9 antibody was a gift from Dr. Robbie Loewith (Urban et al. 2007). Membranes were developed by using Amersham ECL Prime Western Blotting Detection Reagent and visualized by BioRad Gel Doc XR+ gel documentation system. ImageLab software from BioRad was used to crop and quantify the immunoblots.

### Fluorescent microscopy

Cells were grown in YEP-lactate overnight and treated with MMS and rapamycin as described previously (Eapen et al. 2017). Strains that contain mCherry-Atg8 plasmids were grown in leucine drop-out media with 2% dextrose. Cells were harvested by centrifugation at 3000 rpm for 1 min and then washed into synthetic complete media prior to imaging. The images were taken with Nikon Eclipse E600 microscope and processed with Fiji software. FM4-64 dye (Invitrogen) stock solution was prepared and used as previously described (Vida and Emr 1995). Cells containing Atg11, Atg13 and Atg17 foci were counted manually. For each condition, more than 150 cells were quantified.

### Statistical analysis

Statistical analysis was performed by using Prism7 software. Statistical significance was calculated by using 2-way ANOVA, corrected with Dunnett’s comparison test for multiple comparisons. The p values calculated were indicated in the figure legends. The error bars represent standard error.

## ACKNOWLEDGEMENTS

We thank Dr. Eric Baehrecke for his insightful feedback on the manuscript and Dr. Vladimir Denic for useful discussions. This work was supported by NIH grant GM61766.

---

**Supplementary Figure 1.**
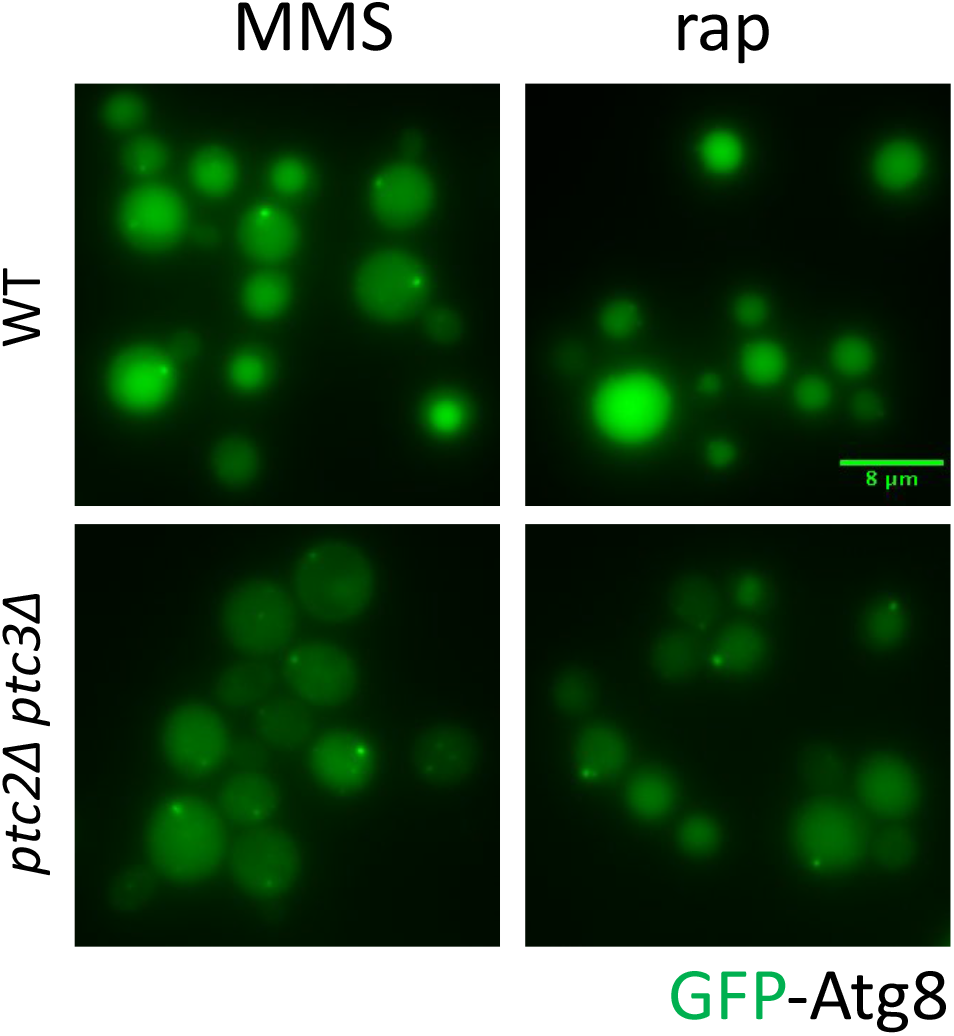
GFP-Atg8 localization in WT and *ptc2Δ ptc3Δ* strains 4h after MMS or rapamycin treatment.

**Supplementary Figure 2.**
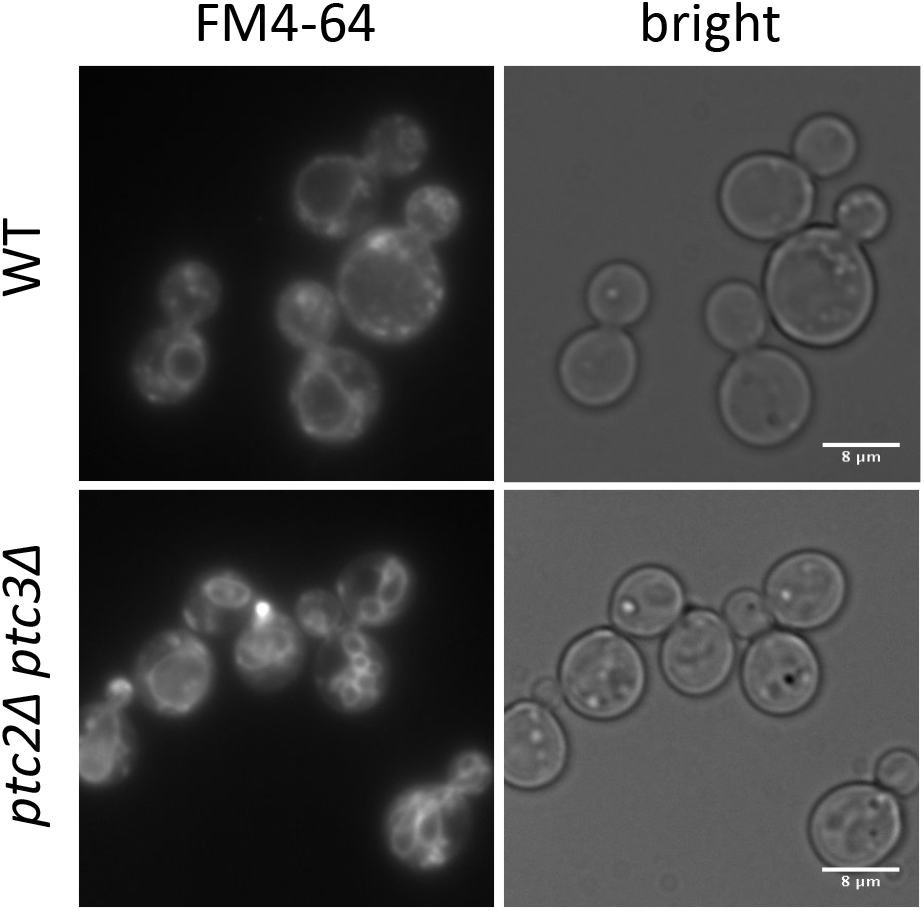
FM4-64 staining of cycling WT and *ptc2Δ ptc3Δ* strains showing vacuolar structures.

**Supplementary Figure 3.**
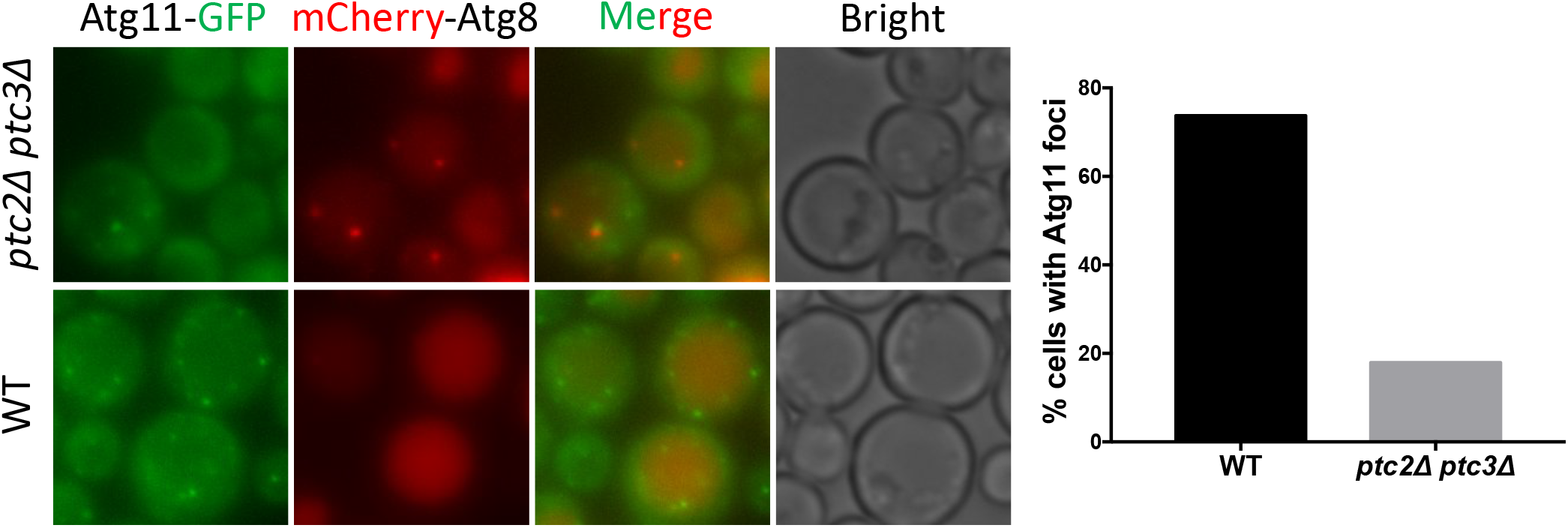
Atg11-GFP and mcherry-Atg8 localization in WT and *ptc2Δ ptc3Δ* mutant strains 4h after rapamycin treatment. Percentage of cells with visible Atg11 foci were counted and plotted.

**Supplementary Figure 4.**
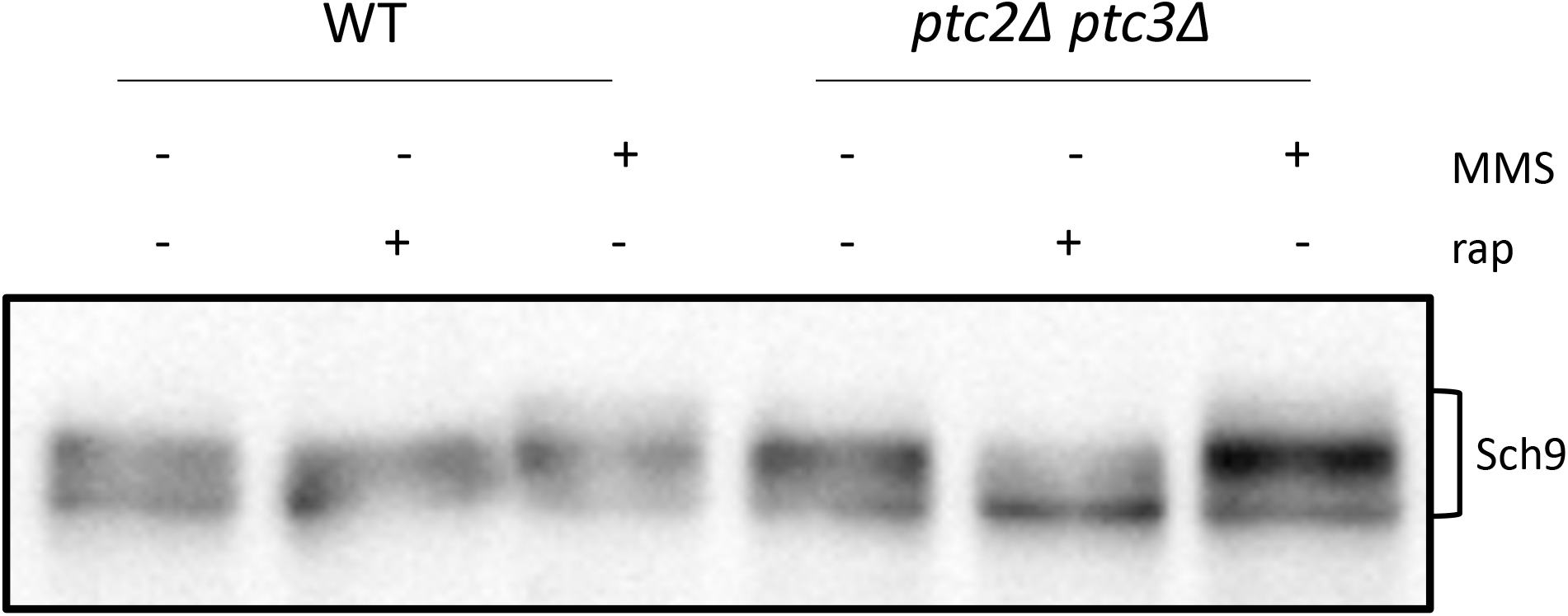
Sch9 phosphorylation in WT and *ptc2Δ ptc3Δ* strains 4h after MMS or rapamycin treatment. Samples were collected as explained before and blotted with anti-Sch9 antibody.

**Supplementary Figure 5.**
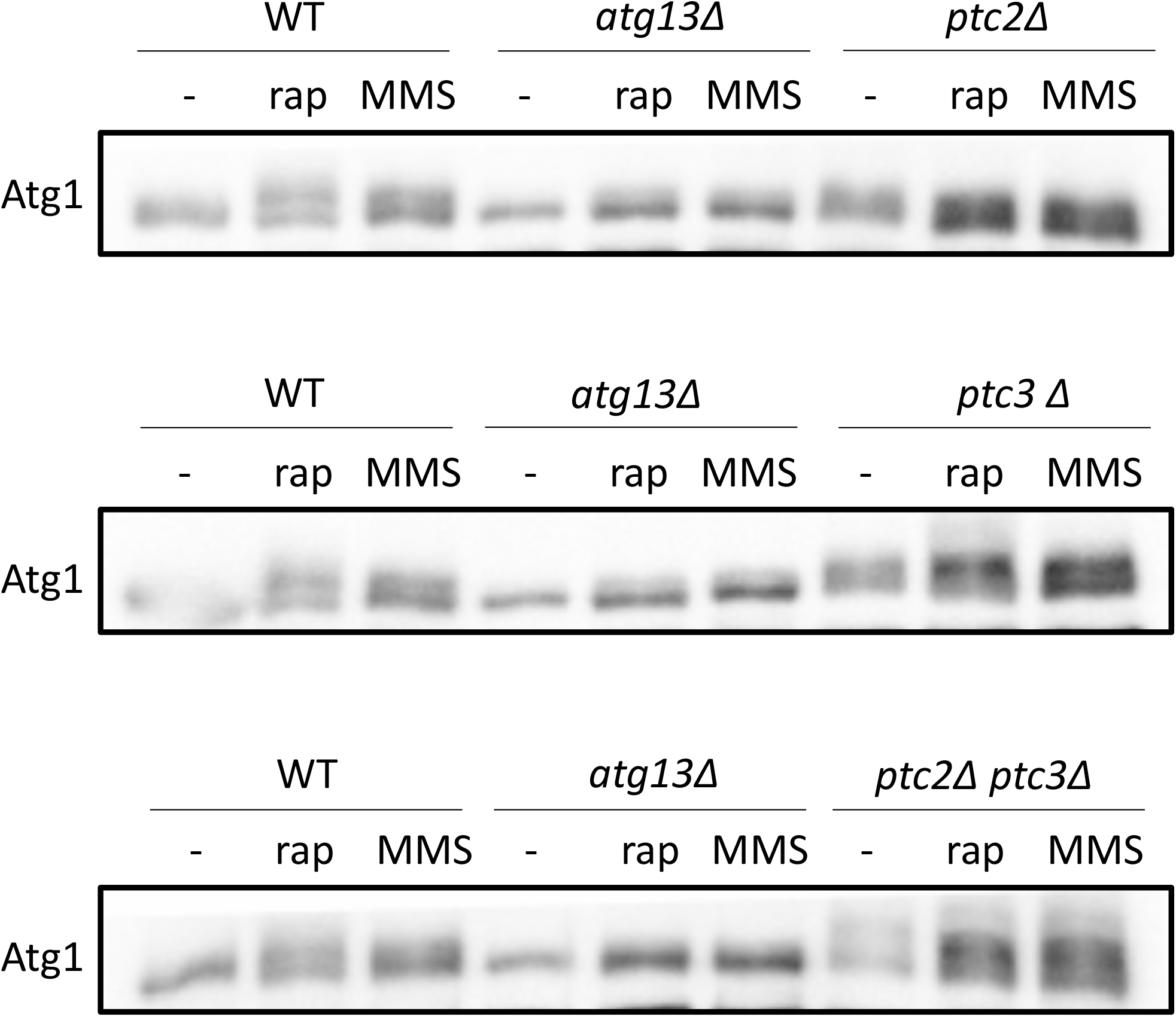
Atg1 hyperphosphorylation in WT, *atg13Δ, ptc2Δ, ptc3Δ* and *ptc2Δ ptc3Δ* strains. Cells were treated with rapamycin and MMS for 4h as previously explained. Samples were collected and analyzed by blotting.

**Supplementary Figure 6.**
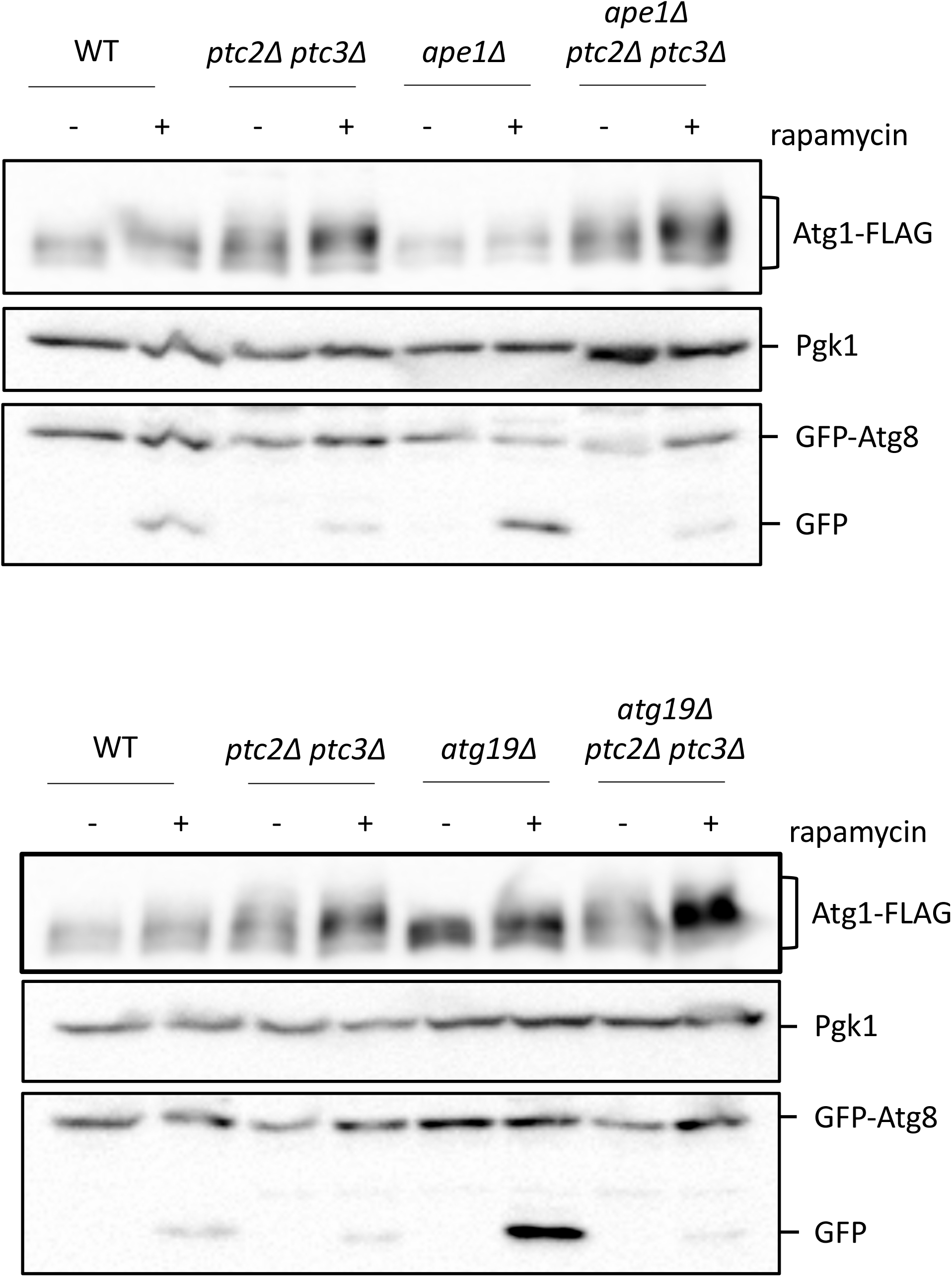
Atg1 hyperphosphorylation in WT, *ape1Δ, atg19Δ, ptc2Δ ptc3Δ, ape1Δ ptc2Δ ptc3Δ* and *atg19Δ ptc2Δ ptc3Δ* mutant strains. Cells were treated with rapamycin as previously explained and blotted for FLAG, Pgk1 and GFP.

**Supplementary Figure 7.**
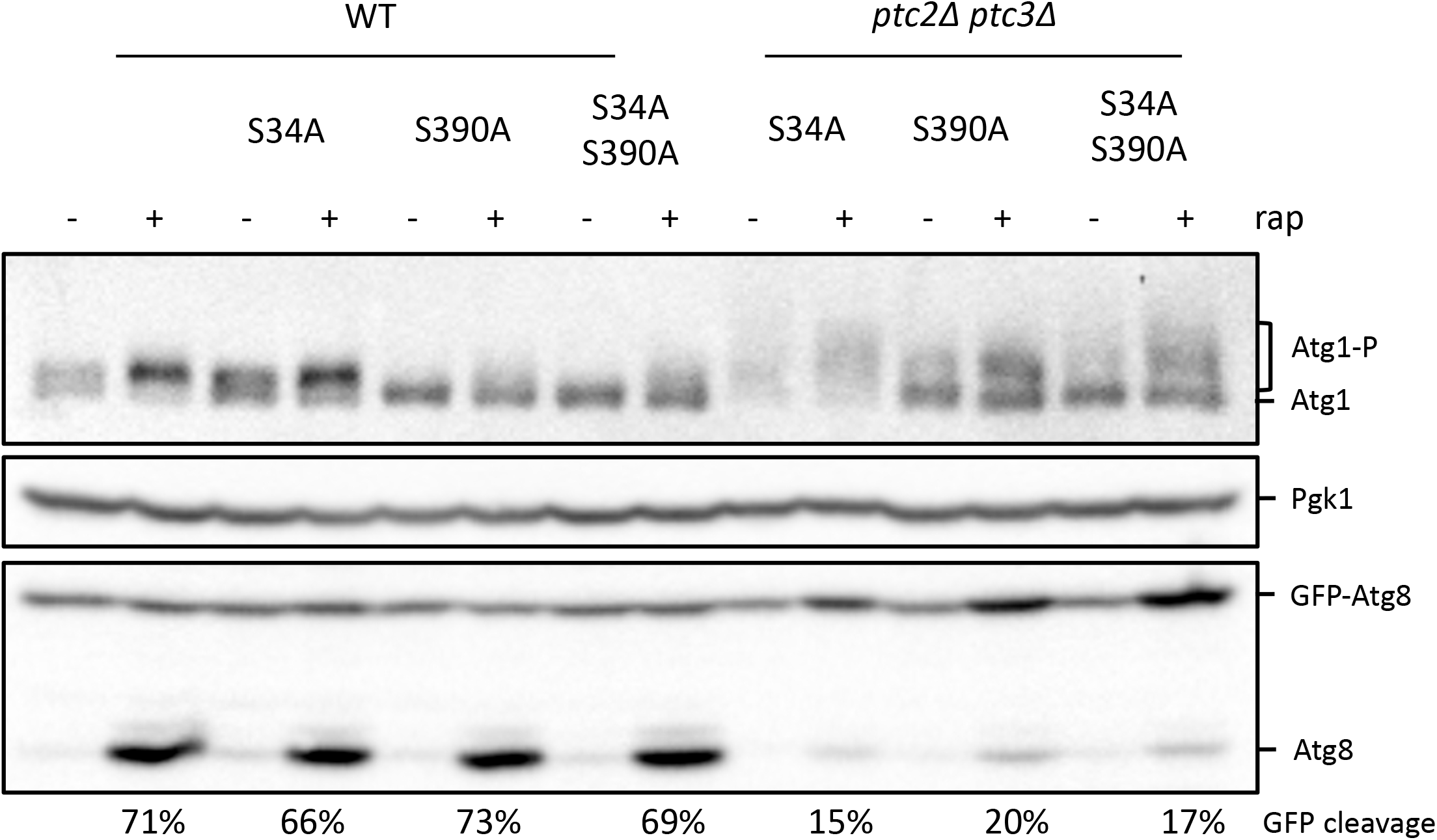
Atg1 hyperphosphorylation in WT, *atg1-S34A, atg1-S390A, atg1-S34AS390A, ptc2Δ ptc3Δ atg1-S34A, ptc2Δ ptc3Δ atg1-S390A* and *ptc2Δ ptc3Δ atg1-S34AS390A* mutant strains 4h after rapamycin treatment. Samples were blotted for Pgk1 as loading control and GFP to measure percent GFP cleavage.

**Supplementary Figure 8.**
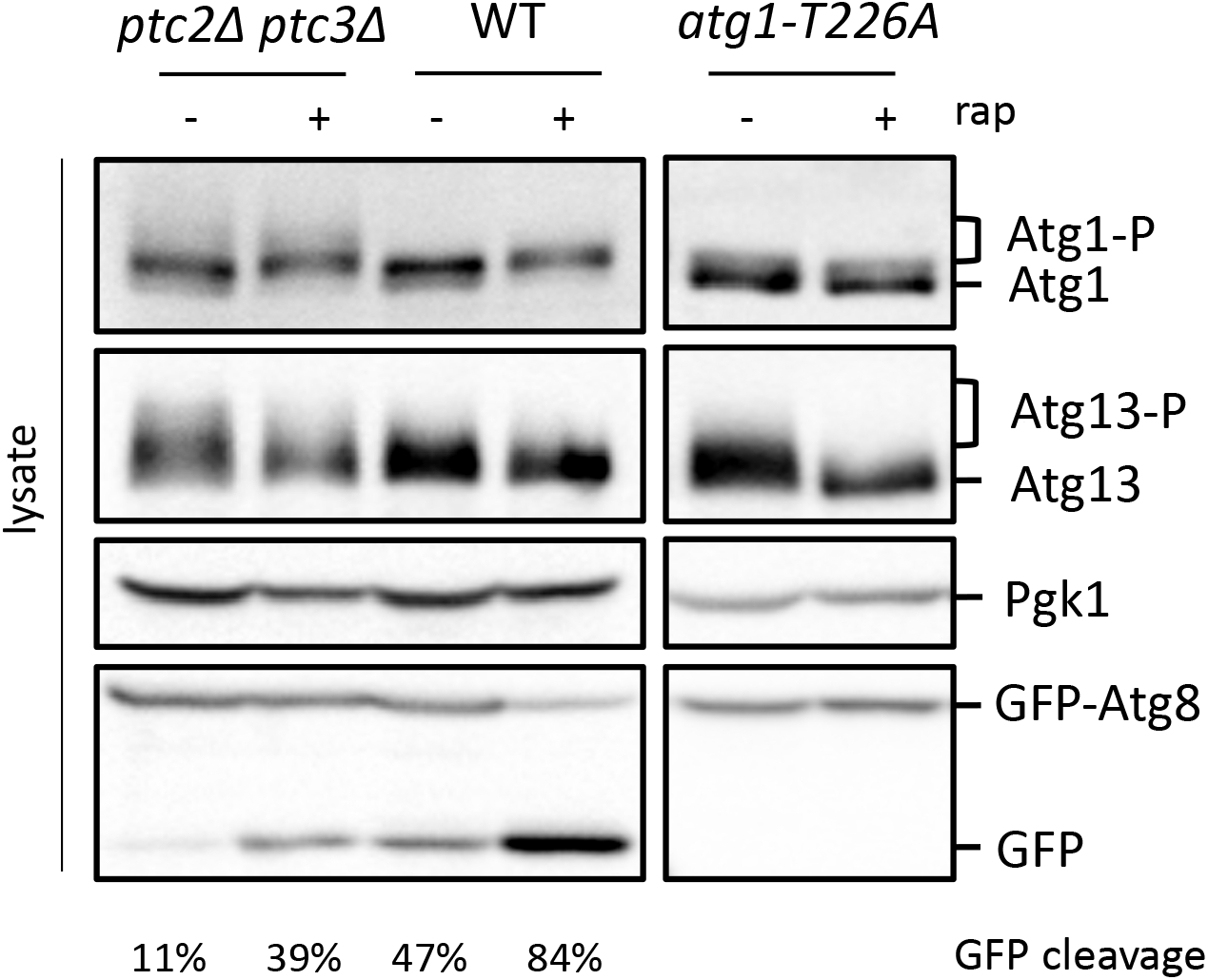
Atg13 and Atg1 hyperphosphorylation in WT, *ptc2Δ ptc3Δ* and *atg1-T226A* mutant strains 4h after rapamycin treatment. Samples were also blotted for Pgk1 for loading control and GFP to measure percent GFP-Atg8 cleavage.

